# Progressive drought transcriptomics and co-expression framework in eggplant (*Solanum melongena* L.)

**DOI:** 10.64898/2026.02.21.706950

**Authors:** Matteo Martina, Cristina Morabito, Andrea Moglia, Anna Maria Milani, Lorenzo Barchi, Alberto Acquadro, Cinzia Comino, Francesca Secchi, Ezio Portis

## Abstract

Drought is a major constraint for eggplant productivity in Mediterranean and semi-arid environments, yet stage-specific molecular features that distinguish tolerant and sensitive genotypes under progressive water deficit remain limited. Here, we profiled drought responses in two contrasting eggplant (Solanum melongena L.) genotypes from the G2P-SOL core collection, integrating physiology and transcriptomics to resolve genotype-dependent programs at moderate and severe stress. Physiological measurements confirmed divergent drought performance, with the tolerant genotype ‘Berenjena de rabo largo’ (GPE020510) maintaining water status and stomatal function longer than the sensitive genotype ‘Qianzi’ (GPE008940). RNA-seq revealed strong transcriptional reprogramming in both genotypes, but with distinct timing and functional priorities across stress transitions. At moderate stress, tolerance was associated with early ABA-centred regulatory control and dehydration protection (including ABI5, TAS14 and LEA/dehydrin-related loci), coupled to transport and redox homeostasis and repression of growth-associated outputs. In contrast, the sensitive genotype showed prominent early regulatory and RNA/protein-turnover signatures alongside weaker representation of cuticle/barrier and chloroplast/light-management functions. Under severe stress, the sensitive genotype shifted toward a broad high-maintenance state enriched in remodeling, detoxification and transporter activity, whereas the tolerant genotype displayed a more targeted adjustment featuring plastid photoprotection, proteostasis and selective metabolic reconfiguration. Co-expression network analysis supported this stage-resolved model by identifying modules and hub genes with contrasting temporal trajectories between genotypes, linking earlier coordinated regulatory/membrane-trafficking and plastid/redox tuning to drought tolerance. Overall, these results indicate that eggplant drought resilience is associated with genotype-specific coordination and timing of protective programs superimposed on a shared basal stress response, and they provide prioritized candidate pathways and genes for functional validation and breeding.

## Introduction

Drought is a dominant constraint for crop productivity and quality (Paladini et al., 2025; Seleiman et al., 2021), and its effects are expected to intensify as climate variability increases the frequency of irregular rainfall and heat–dry spells (Abbass et al., 2022; Araus and Kefauver, 2018; Dai, 2013; Haile et al., 2024). For fleshy-fruited vegetable crops, water deficit is particularly impactful because yield formation relies on sustained carbon assimilation, source–sink coordination, and maintenance of cellular homeostasis during prolonged developmental windows (Ahluwalia et al., 2021; Fahad et al., 2017; Seleiman et al., 2021; Yıldız and Akı, 2025). In Solanaceae, drought commonly triggers a cascade of interconnected processes such as stomatal limitation and altered carbon allocation, membrane and cell-wall remodeling, shifts in redox balance and detox capacity, and a progressive reconfiguration of growth programs that together determine whether plants remain functional under moderate stress or transition to severe-stress states that can be associated with irreversible performance losses (Blanchard-Gros et al., 2022; Emami Bistgani et al., 2024; Maioli et al., 2024; Martina et al., 2024; Ojeilua et al., 2025; Pang et al., 2024; Saptiningsih et al., 2023; Wadood et al., 2024; Yıldız and Akı, 2025). As drought typically develops along a continuum rather than as a binary event, a key conceptual challenge is to understand how plants move from moderate to severe stress and which molecular features distinguish these stages.

Eggplant (*Solanum melongena* L.) is a major vegetable crop in Mediterranean and semi-arid regions, where tightening irrigation and more frequent heat–drought episodes increasingly penalize yield and quality (Gaccione et al., 2023; Sękara et al., 2016; Toppino et al., 2022; Villanueva et al., 2026). Yet, compared with model species, drought-response frameworks in eggplant are still relatively fragmented, and markers that robustly separate tolerant from susceptible genotypes remain limited. Recent studies nonetheless provide clear mechanistic and genomic entry points. Functional evidence shows that the bHLH regulator SmMYC2 can enhance drought resilience by promoting secondary cell wall thickening through SmNST1-linked regulation, supporting cell-wall reinforcement as a relevant tolerance output(Li et al., 2025). Severity-aware transcriptomics further indicate that drought responses depend strongly on stress intensity and developmental context: RNA-seq across two PEG levels and two exposure times showed that the wild relative (*S. dasyphyllum*) is more water-deficit tolerant than cultivated eggplant and deploys broader drought TF repertoires (AP2/ERF, DREB, bZIP, WRKY, bHLH), with ABA signaling as a central axis and additional involvement of phenylpropanoid/flavonoid and carotenoid metabolism, chlorophyll metabolism, and photosynthesis-related pathways under both moderate and severe regimes (Villanueva et al., 2023). Physiological studies across wild relatives and interspecific hybrids similarly report strong drought impacts on growth and tissue water content together with increases in biochemical stress markers such as proline and MDA, underscoring polygenic control and diverse stress strategies (González-Orenga et al., 2023). However, single time points or single severity levels can confound early acclimation with late damage-control programs. Disentangling these phases requires designs that explicitly compare moderate and severe drought in contrasting genotypes and interpret expression as stage-specific programs rather than static lists of “drought genes”.

Here, we address this gap by contrasting two eggplant genotypes with divergent drought performance across progressive stress stages, focusing on transitions that capture (i) the onset of moderate drought (T0 to WS-M) and (ii) drought escalation from moderate to severe stress (WS-M to WS-S). We integrate three complementary transcriptome layers-differential expression, GSEA-based functional directionality, and WGCNA module/hub structure to ask whether tolerance is associated with early engagement of protective and homeostasis-buffering programs, and whether susceptibility is associated with early regulatory rewiring that is not accompanied by coherent deployment of barrier- and plastid-support functions. We further test, at the level of co-expression architecture, whether the WS-M to WS-S transition is dominated by broad severity-associated “maintenance-mode” programs in the susceptible genotype, versus more targeted plastid/light-management and redox-associated programs in the tolerant genotype.

By building a staged, genotype-aware framework, this study aims to move beyond single-gene narratives and instead provide a systems description of drought-response trajectories in eggplant. The resulting interpretation yields (i) stage-resolved candidate processes that can be linked to tolerance versus susceptibility as hypotheses, (ii) prioritized candidate genes and module hubs that anchor these hypotheses, and (iii) a template for future targeted experimentation to test whether the inferred programs translate into measurable protective outputs.

## Materials and Methods

### Plant material and experimental design

Two eggplant (*Solanum melongena* L.) genotypes from the G2P-SOL core collection (http://www.g2p-sol.eu/) were selected based on their contrasting responses to drought stress previously characterized within the G2P-SOL framework. The drought-tolerant genotype ‘*Berenjena de rabo largo*’ (GPE020510), originating from Spain, and the drought-sensitive genotype Qianzi (GPE008940), originating from China, were used in this study. Seeds were germinated in a growth chamber under controlled conditions (25/18 °C day/night, 16 h light/8 h dark photoperiod, 60–70% relative humidity). At the four-leaf stage, seedlings were transplanted into 1 L pots filled with a standardized peat/perlite substrate (3:1, v/v) and transferred to a greenhouse for further growth. Thirty-eight plants were irrigated daily to pot capacity until the onset of drought treatment. At the 12-leaf stage, 18 plants (9 per genotype) were maintained under well-watered conditions (controls), while the remaining 20 (10 for genotype) were subjected to a progressive reduction in water availability (stressed) starting on day 0. Water stress was imposed by restoring 80% of the water lost on the previous day, with pot weight measured every morning. Drought stress was imposed gradually and monitored through plant water status, allowing the definition of two physiological sampling points: moderate water stress (WS-M), corresponding to a leaf water potential of approximately −1.0 MPa, and severe water stress (WS-S), corresponding to a leaf water potential of approximately −2.0 MPa. Plant sampling (n=3 for controls, n=5 for stressed) for physiological and transcriptomic analyses was performed at these defined physiological stages.

### Physiological measures

Stem water potential (Ψ) was measured during drought progression. Measurements were performed at midday on fully expanded, non-transpiring leaves using a Scholander-type pressure chamber, following standard procedures (Morabito et al., 2022). For each genotype and condition, three individual plants were selected and one leaf per plant was measured. Stem water potential data were used to quantify the severity of water stress and to compare the physiological responses of the two eggplant genotypes under control and drought conditions.

Stomatal conductance was measured on fully expanded leaves using a portable gas exchange system (LI-6800, LI-COR Biosciences). CO_2_ concentration entering the cuvette was set to 400 µmol CO_2_ mol^−1^ air, and relative humidity was set to follow environmental conditions. Measurements were conducted on all plants (one leaf per plant) between 10.00 a.m. and 12.00 p.m. under ambient environmental conditions to minimize diurnal variation.

### RNA extraction and sequencing

For transcriptome profiling, fully expanded leaves were collected at each sampling point from control and stressed plants of the two eggplant genotypes. Sampling was carried out on the same plants and at the same time point used for leaf water potential measurements. For each genotype and treatment, three biological replicates were collected, each consisting of a pooled leaves sample from an individual plant, which was immediately frozen in liquid nitrogen and stored at −80°C until processing. Total RNA was extracted from approximately 100 mg of powdered leaf tissue using a column-based plant RNA purification kit (Spectrum™ Plant Total RNA Kit, Sigma-Aldrich), followed by DNase treatment to remove residual genomic DNA. RNA concentration and purity were assessed spectrophotometrically, and RNA integrity was verified by agarose gel electrophoresis and capillary electrophoresis. Stranded mRNA libraries were prepared from poly(A)-enriched RNA according to standard Illumina protocols and sequenced as paired-end reads (2 × 150 bp) on an Illumina NovaSeq X Plus at Novogene Europe (Cambridge,UK) and Igatech (Udine, IT).

### Read processing, transcript quantification and differential expression analysis

Raw sequencing reads were quality filtered and adapter-trimmed using fastp. Cleaned reads were pseudo-aligned to the *Solanum melongena* reference transcriptome (SMEL v5, ‘67/3’ line; Gaccione et al., 2025) using Salmon in quasi-mapping mode with default parameters. Transcript abundances were quantified as counts and Transcripts Per Million (TPM), and gene-level count matrices were imported into R (Patro et al., 2017) using the tximport package (Soneson et al., 2015). Differential expression analysis was performed using DESeq2 (Love et al., 2014 - design *geno + condition + geno:condition*). The *geno* factor included two levels (GPE008940 and GPE020510), while *condition* represented the combination of time point (T0, WS-M and WS-S) and treatment (CTR and STR), with T0_CTR set as the reference level. Size-factor normalisation and dispersion estimation followed the standard DESeq2 workflow, and differential expression was assessed using Wald tests. Biologically relevant contrasts were defined to compare drought-stressed and control plants within each physiological stage, to evaluate transcriptional changes along drought progression, and to compare genotypes under identical experimental conditions. *p-values* were adjusted for multiple testing using the Benjamini–Hochberg false discovery rate (FDR - Benjamini and Hochberg, 1995). Genes with adjusted p-value < 0.05 were considered differentially expressed, and an additional threshold was applied for selected visualisations (|log₂ fold change| > 1). For exploratory analyses, size-factor normalised counts were transformed using a log₂(count + 1) transformation and used for principal component analysis and heatmap generation with pheatmap (Kolde, 2010).

### Gene set enrichment analysis (GSEA)

To investigate transcriptional changes at the pathway level, we performed Gene Set Enrichment Analysis (GSEA) using (Langfelder and Horvath, 2008) the *enrichplot* package (https://github.com/YuLab-SMU/enrichplot) in R. For each contrast of interest, all genes with non-missing log₂ fold change were ranked according to the DESeq2 log₂ fold change, with positive values indicating higher expression in the second level of the contrast. These ranked gene lists were used as input for the *gseGO* function, employing a dedicated *S. melongena* annotation package and SMEL5 gene identifiers as *keys*. Gene Ontology enrichment was carried out separately for the Biological Process, Molecular Function and Cellular Component ontologies, using a minimum gene set size of 5 and a maximum of 500 genes. Significance of enrichment scores was assessed using a simple permutation scheme with 20,000 permutations, and *p-values* were adjusted for multiple testing using the Benjamini–Hochberg procedure. GO terms with an adjusted *p-value* ≤ 0.05 were considered significantly enriched. Enriched GO categories were visualized using the *enrichplot* package. Ridgeplots were generated to display, for each GO term, the distribution of ranked log₂ fold changes of core enriched genes, with color scales encoding the adjusted p-value. In addition, mirrored dot plots were constructed to compare enrichment patterns between the two genotypes by jointly displaying, for each GO term, its presence in GPE022290 and/or GPE036890, and the size of the corresponding core gene sets. This representation allowed us to distinguish terms that were uniquely enriched in the sensitive line, uniquely enriched in the tolerant line, or shared between them.

### Co-expression network analysis (WGCNA)

Weighted gene co-expression network analysis (WGCNA) was performed in R using the WGCNA package (Langfelder and Horvath, 2008) to identify modules of co-expressed genes showing coordinated transcriptional dynamics during drought progression in the two eggplant genotypes. Genes with low expression across samples were filtered prior to network construction. Normalized expression values derived from DESeq2 size-factor normalization were used as input and transformed to stabilize variance. To focus on transcriptional changes associated with stress progression rather than absolute baseline differences between genotypes, the analysis was conducted on expression changes relative to the baseline condition. For each genotype, expression values were centered by subtracting the genotype-specific mean expression at the T0_CTR condition, generating Δ-expression values that capture stage-dependent transcriptional variation across subsequent physiological stages. Pairwise gene co-expression similarity was quantified using Pearson correlations on Δ-expression values, and an adjacency matrix was constructed by applying a soft-thresholding power selected according to the scale-free topology criterion. The adjacency matrix was transformed into a topological overlap matrix (TOM), and genes were hierarchically clustered based on TOM dissimilarity. Co-expression modules were detected using dynamic tree cutting and subsequently merged based on eigengene similarity. Module eigengenes, defined as the first principal component of each module, were used to summarize module-level expression patterns across samples and to evaluate genotype-dependent trajectories along drought progression. Hub genes were identified within selected modules based on intramodular connectivity and module membership (kME) values.

## Results

### Physiological responses to water stress differ between genotypes

Under well-watered conditions (day 0), no significant differences were observed between the two genotypes (*i.e.* GPE008940 and GPE020510) for either stomatal conductance or stem water potential, indicating comparable physiological status at the onset of the experiment (Fig. 1). As water stress progressed, clear genotype-dependent differences emerged. Under water-limited conditions, stomatal conductance declined in both genotypes, but with markedly different dynamics. In GPE008940, stomatal conductance (gs) decreased sharply after 5 days of treatment, and at that time the stem water potential (ψstem) dropped to approximately -1MPa. Two days later, this genotype closed its stomata, and the ψstem reached values of about -2MPa. In contrast, GPE020510 did not show changes in either gs or ψstem after 5 days of stress imposition compared to their controls. These plants required more days to reach moderate and severe stress levels comparable to those experienced by the other genotype.

**Figure 1.**
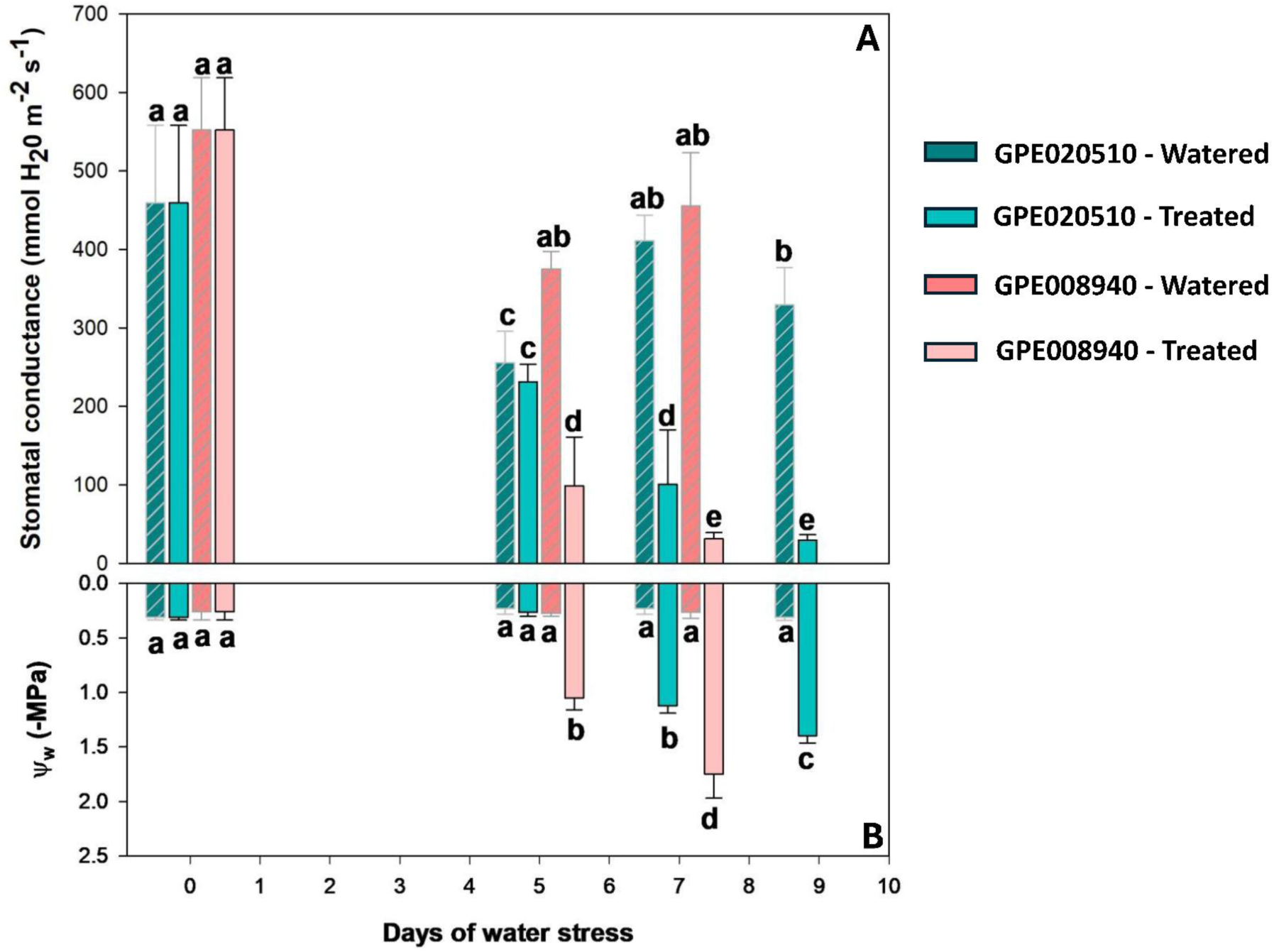
(A) Stomatal conductance (gs; mmol H₂O m⁻² s⁻¹) and (B) leaf water potential (Ψw; MPa) measured over the course of increasing days of water deficit. Bars represent the two genotypes (GPE020510 - green - and GPE008960 - red) under well-watered and water-stressed conditions (as indicated by the bar color/pattern in the legend). Data are shown as mean ± SE. Different letters above bars denote statistically significant differences among treatments/genotypes within each time point (post hoc test, P < 0.05).

In GPE020510 plants, the stomatal conductance decreased after 7 days of stress, and the stomata were fully closed after 9 days, when stem water potential dropped to approximately -1.5 MPa.

Under well-watered conditions, both genotypes maintained stable stomatal conductance and stem water potential throughout the experiment, confirming that the observed differences were specifically associated with the water stress treatment rather than with developmental effects.

### Transcriptomic variation under stress conditions

To assess genotype-dependent transcriptional divergence under drought stress, differential expression analyses were performed between GPE008940 and GPE020510 at two stress conditions, moderate (WS-M) and severe (WS-S). Principal component analysis revealed that, while drought stress accounted for the major source of transcriptomic variation, clear genotype-specific separation was evident within both stress conditions, with distinct patterns emerging at WS-M and WS-S (Figure 2A). Consistently, pairwise comparisons between genotypes identified thousands of differentially expressed genes under both stress levels (Figure 2B). As depicted by upset plot (Figure 2C), a core set of 1032 genes was shared by the two genotypes between WS-M and WS-S, together with stress-severity-specific gene subsets (81 in WS-M and 1043 in WS-S), suggesting that drought intensity modulates the transcriptional programs underlying genotype-dependent responses.

**Figure 2.**
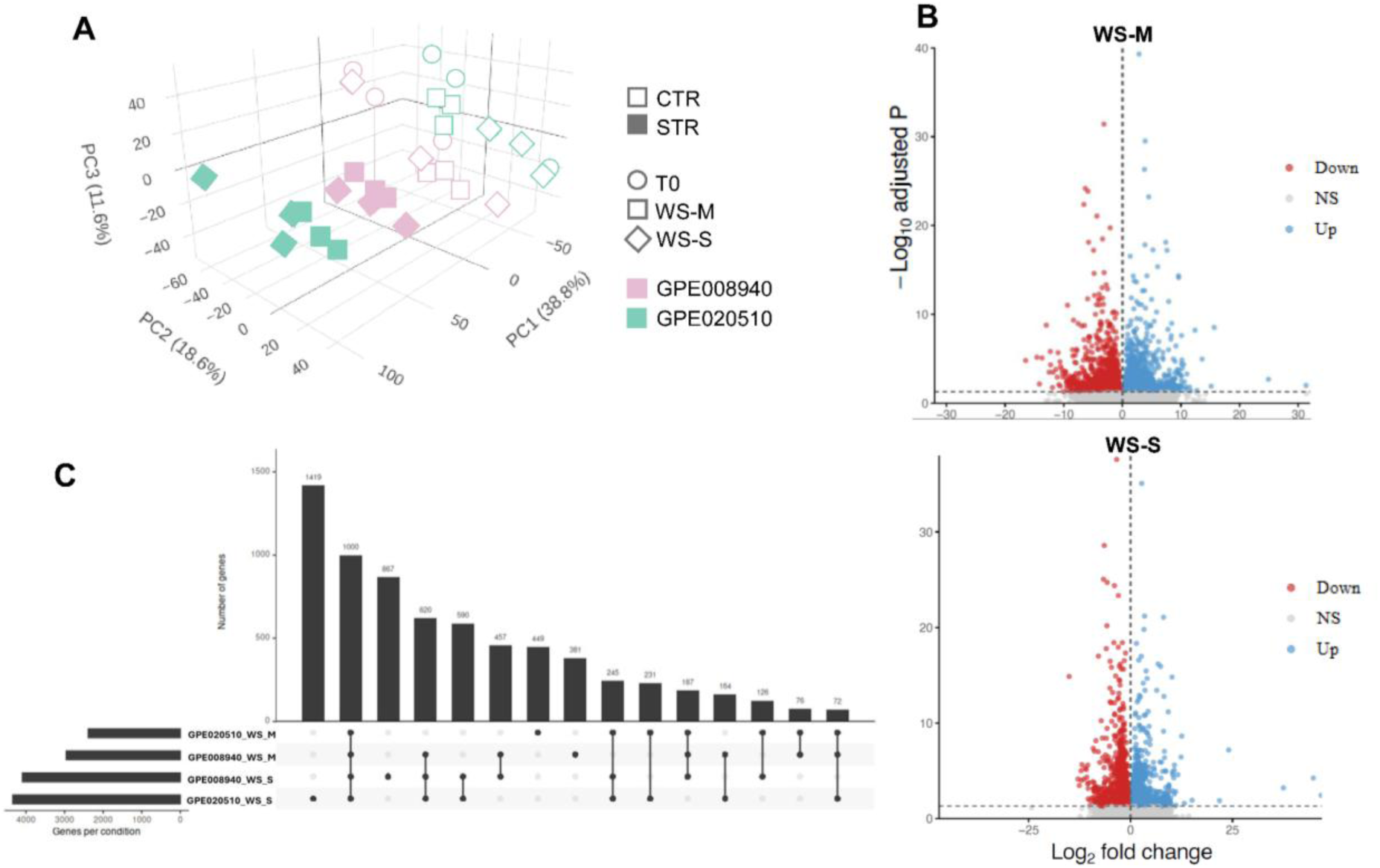
(A) Three-dimensional PCA of variance-stabilized RNA-seq expression profiles showing clustering of samples by genotype and drought severity (moderate, WS-M; severe, WS-S), indicating progressive transcriptome reprogramming with stress intensification. Axes report the percentage of variance explained by each principal component. (B) Volcano plots of differential expression highlighting drought-responsive genes (FDR-adjusted P and log₂ fold-change thresholds as indicated by dashed lines), with upregulated genes in blue, downregulated genes in red, and non-significant genes in grey. (C) UpSet plot summarizing the size and overlap of drought-responsive gene sets across genotype × severity contrasts, illustrating shared and genotype-/stage-specific transcriptional programs.

At both moderate (WS-M) and severe (WS-S) drought stress conditions, differentially expressed genes were distributed across a broad set of Gene Ontology categories, indicating extensive transcriptional remodeling under water deficit (Figure 3).

**Figure 3.**
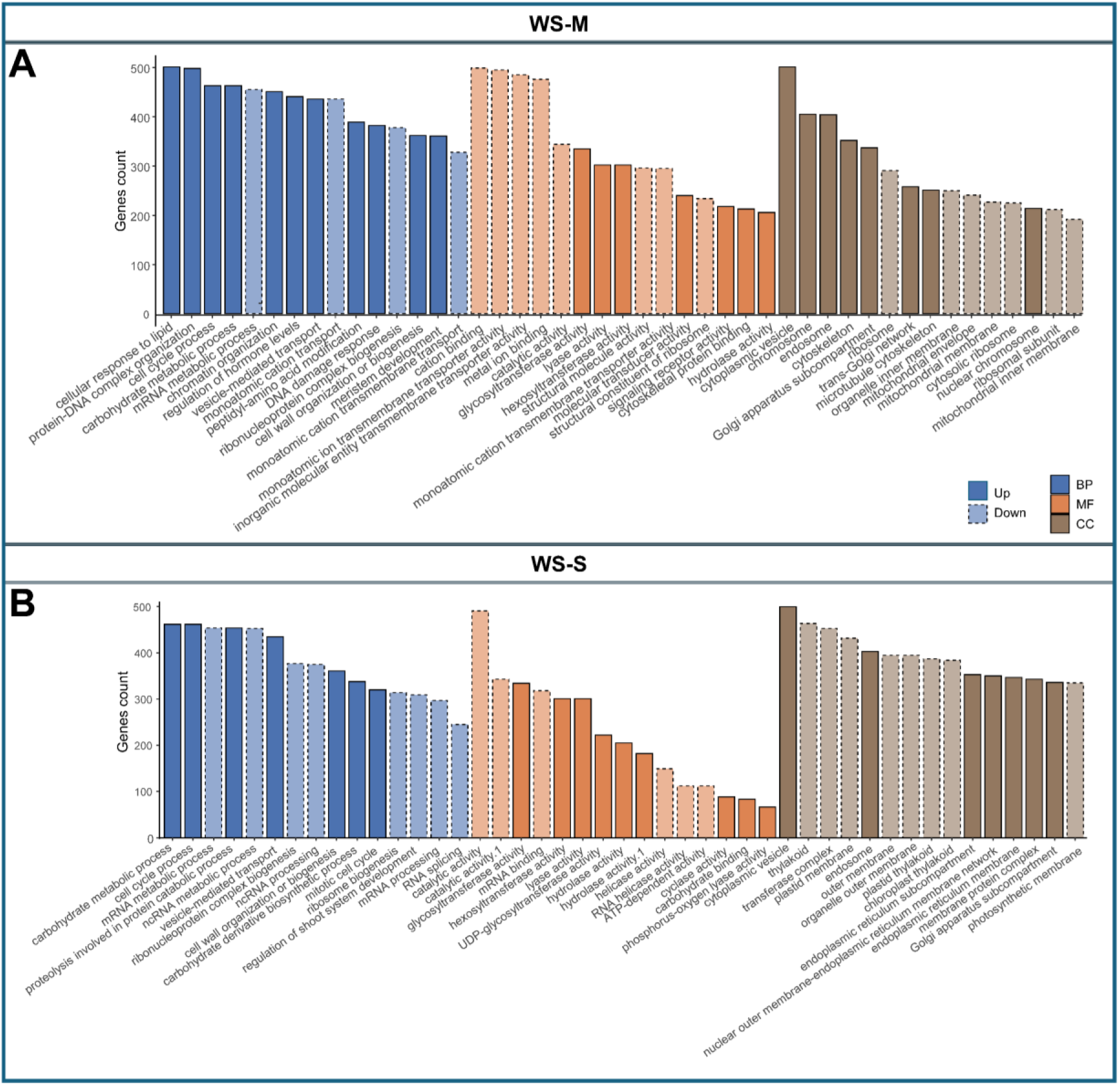
Bar plots summarize significantly enriched Gene Ontology (GO) terms among differentially expressed genes, separated by GO domain (BP, biological process; MF, molecular function; CC, cellular component) and regulation direction (Up vs Down, as indicated by solid vs dashed bars). The upper and lower panels correspond to the two drought transitions/severity contrasts analyzed (moderate - A - and severe drought - B), highlighting a shift from early regulatory/transport-associated responses to stronger repression of photosynthesis- and growth-related categories and enhanced stress/maintenance-associated functions as drought severity increases.

### Susceptible genotype shows early regulatory rewiring and late stress escalation under increasing drought severity

At the transition from well-watered conditions to moderate drought (T0 to WS-M), the susceptible genotype GPE008940 showed pronounced transcriptional changes (5328 upregulated and 5852 downregulated genes, Figure 4C). Several strongly induced transcripts were associated with drought-and stress-related functions, including an abscisic acid and environmental stress–inducible protein (TAS14), low-temperature-induced proteins, heat shock proteins, and multiple transport-related genes such as aquaporins and sodium-coupled amino acid transporters. In parallel, transcriptional regulators and signaling components, including NAC and homeobox–leucine zipper transcription factors, were also upregulated. Conversely, a subset of genes related to cell wall components, hormone-regulated proteins, and receptor-like kinases showed significant downregulation at this stage.

**Figure 4.**
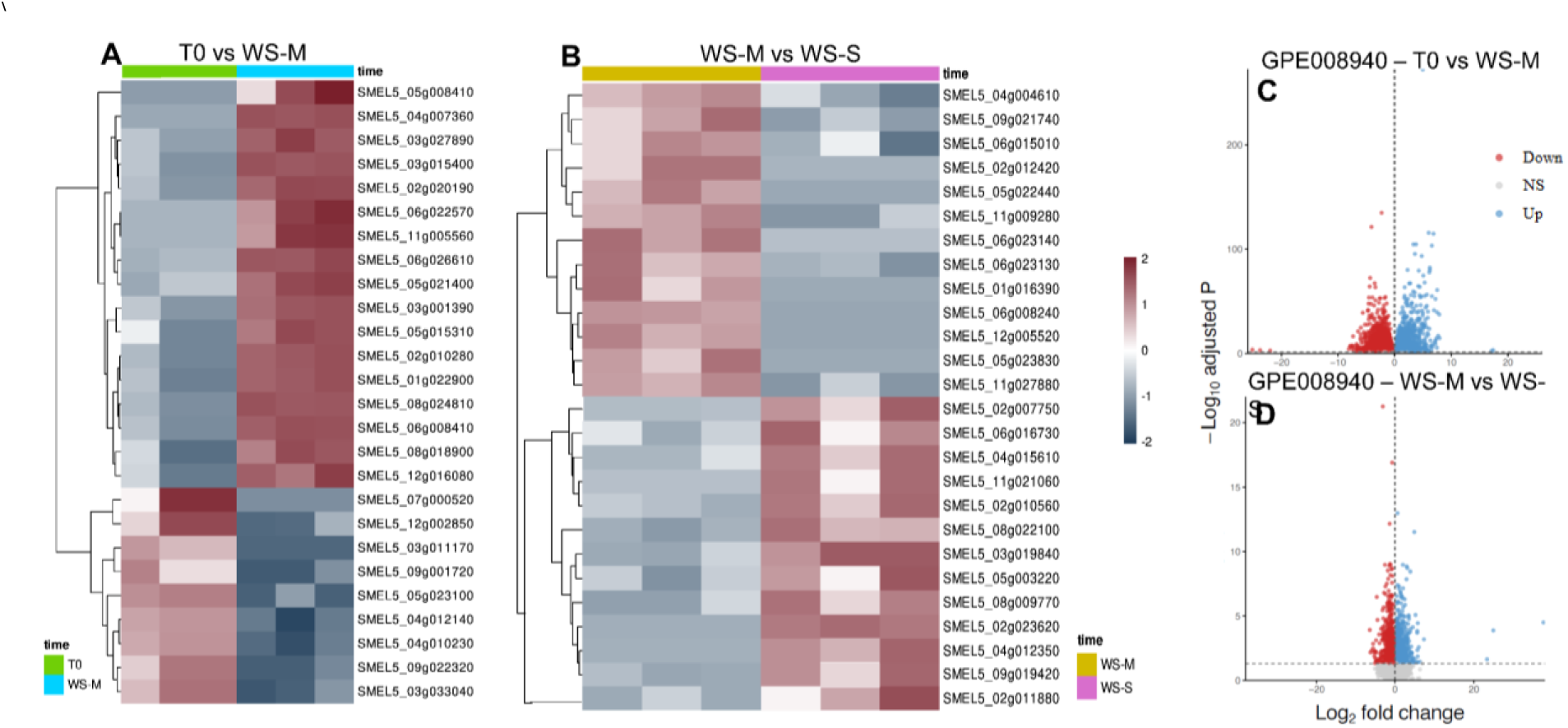
Heatmaps (left) show scaled expression patterns of representative drought-responsive genes across samples, grouped by genotype and drought severity (A - WS-M and WS-S - B; color bars indicate sample classes). Genes are clustered by similarity, highlighting genotype- and stage-dependent transcriptional trajectories. Volcano plots (right; C–D) summarize differential expression for the corresponding contrasts, with significantly upregulated genes shown in blue, downregulated genes in red, and non-significant genes in grey (dashed lines indicate the applied fold-change and adjusted P-value thresholds).

During the subsequent transition from moderate to severe drought (WS-M to WS-S), the transcriptional profile shifted toward genes associated with tissue remodeling and late stress responses (1215 upregulated and 1234 downregulated, Figure 4D). Differentially expressed genes included senescence-associated proteases, chitinases, expansins, extensins, and glycine-rich cell wall proteins, indicating extensive modulation of cell wall–related processes. Several enzymes involved in secondary metabolism and redox regulation, such as laccases, dioxygenases, and glutaredoxins, were also differentially expressed. In addition, genes encoding sugar transporters of the SWEET family, hormone-related enzymes, and RNA-binding proteins showed significant regulation, reflecting further reorganization of metabolic and regulatory processes as drought severity increased.

Gene set enrichment analysis (GSEA, Figure 5) revealed a marked functional separation between activated and suppressed gene sets during the early drought stage (T0 to WS-M; Figure 4A,B). Among activated biological processes, enrichment was dominated by intracellular transport and localization categories (intracellular protein transport, protein transport, establishment of protein localization), together with processes associated with protein–DNA complex organization and mRNA metabolic process. This pattern was supported by the induction of representative nuclear/RNA-processing components, including a U11/U12 snRNP 35 kDa component (SMEL5_09g024500) and a nuclear export factor SDE5 (SMEL5_07g019890; Table 1). In addition, proteasome- and peptidase-related terms (proteasome complex, peptidase complex) were enriched among activated gene sets, consistent with concurrent regulation of protein turnover pathways; among the high-amplitude DEGs, an EID1-like F-box protein (SMEL5_06g029750; Table 1) exemplified this component of the response. In parallel, a subset of the largest DEGs included regulators linked to light/clock and growth-associated signaling, including LUX (SMEL5_06g005050), a PIF1-like factor (SMEL5_06g002810), and GA2-oxidase (SMEL5_04g003260; Table 1).

**Figure 5.**
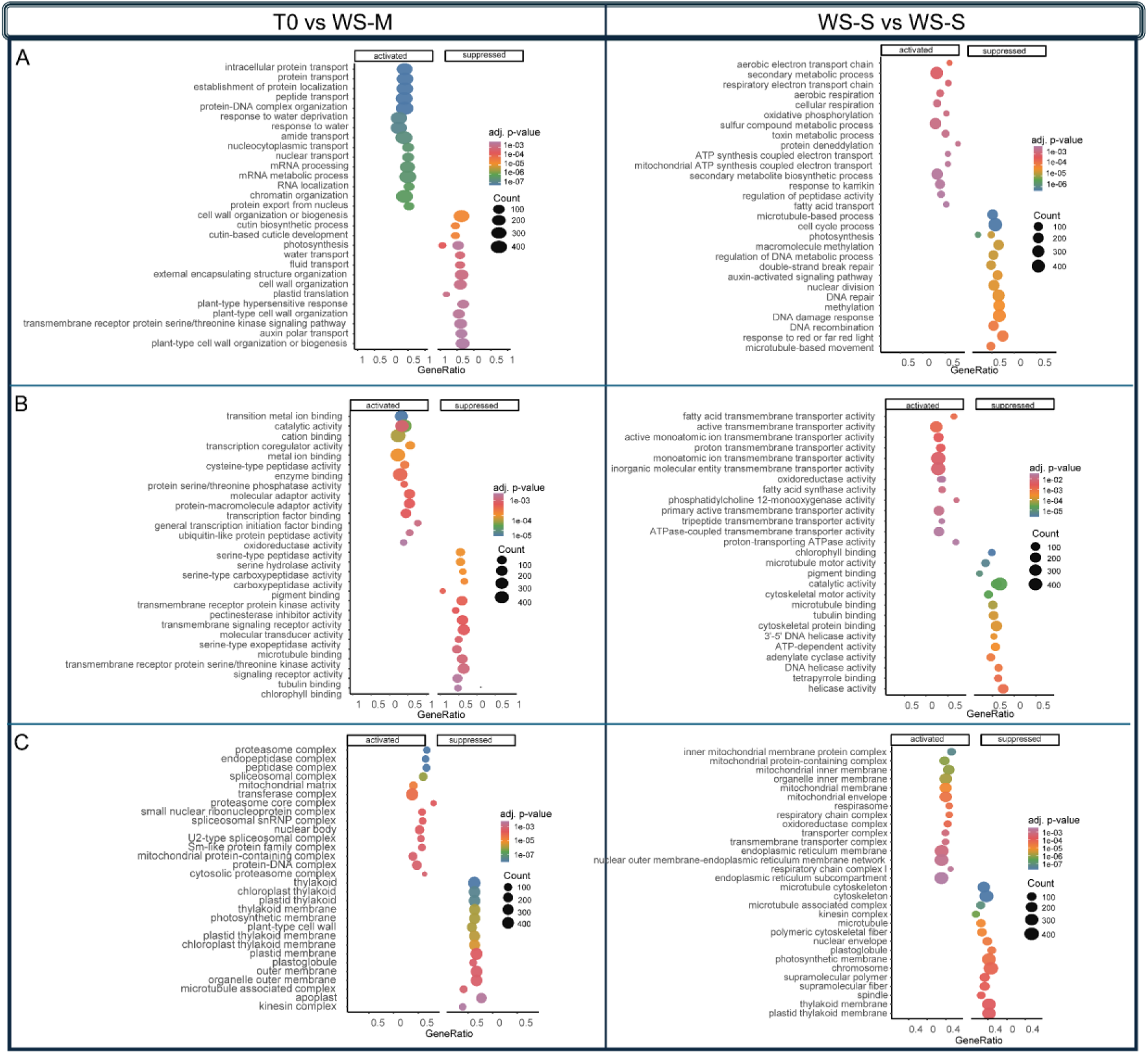
Dot plots summarize significantly enriched terms among upregulated (“activated”) and downregulated (“suppressed”) gene sets for each genotype and drought transition (T0 to WS-M and WS-M to WS-S). Panels report enrichment for Biological Process (A), Molecular Function (B), and Cellular Component (C), highlighting coordinated early regulatory and transport-related activation at moderate stress and a stronger suppression of photosynthesis/light-harvesting and growth-associated functions under severe stress, together with activation of mitochondrial respiration, detoxification, and membrane/lipid remodeling pathways. Dot size reflects gene-set size (count) and dot colour indicates adjusted P value.

**Table 1.** Key drought-responsive genes prioritized in the susceptible genotype GPE008940 across progressive drought severity. The table reports the selected candidate transcripts (gene IDs) highlighted in the Results for the “T0 to WS-M” and “WS-M to WS-S” transitions, with functional annotation, direction of regulation in each contrast, and gene ontology.

| Transition | Gene ID | Function | log2FC | GO |
| --- | --- | --- | --- | --- |
| T0 to WS-M | SMEL5_07g019890 | Nuclear export factor SDE5 | 1.05 | ta-siRNA processing |
|  | SMEL5_06g029750 | EID1-like F-box protein | 1.72 | leaf development |
|  | SMEL5_10g022180 | TIP1-1-like aquaporin | 2.67 | leaf development |
|  | SMEL5_04g003260 | GA2-oxidase | 3.04 | gibberellin biosynthetic process |
|  | SMEL5_06g002810 | PIF1-like factor | 3.34 | gibberellin biosynthetic process |
|  | SMEL5_06g005050 | LUX | 3.59 | regulation of circadian rhythm |
|  | SMEL5_09g024500 | U11/U12 snRNP 35 kDa component | 3.64 | mRNA splicing, via spliceosome |
|  | SMEL5_07g000910 | CER3 | -1.03 | wax biosynthetic process |
|  | SMEL5_01g010820 | SOQ1 | -1.26 | metal ion binding |
|  | SMEL5_01g010830 | SOQ1 | -1.27 | metal ion binding |
|  | SMEL5_06g014400 | TIP-type RB7-5A aquaporin | -1.32 | water transport |
|  | SMEL5_04g002050 | NDH lumenal subunit | -1.43 | photosystem I assembly |
|  | SMEL5_11g014820 | Plastid RNA polymerase β' | -1.49 | DNA-templated transcription |
|  | SMEL5_05g012270 | NIP2-1-like aquaporin | -1.55 | channel activity |
|  | SMEL5_11g028180 | GPAT5-like | -1.66 | cutin biosynthetic process |
|  | SMEL5_01g030260 | Wax ester synthase / DGAT11-like | -1.73 | triglyceride biosynthetic process |
|  | SMEL5_09g018400 | ELIP1-like | -2.79 | photosynthesis |
|  | SMEL5_01g006970 | ABCG transporter | -3.06 | response to abscisic acid |
|  | SMEL5_04g005030 | MYB41-like regulator | -3.82 | response to abscisic acid |
| WS-M to WS-S | SMEL5_02g006090 | PLA1-II-like phospholipase | 1.68 | phospholipid metabolic process |
|  | SMEL5_04g010990 | Metallothionein-like type 2 | 2.3 | metal ion binding |
|  | SMEL5_01g015930 | PLIP1 phospholipase A1 | 2.4 | triglyceride biosynthetic process |
|  | SMEL5_09g022110 | PDR1-like ABC exporter | 2.43 | hormone transport |
|  | SMEL5_07g019280 | ABCG10-like exporter | 2.57 | ABC-type transporter activity |
|  | SMEL5_03g032040 | NADH-quinone oxidoreductase subunit | 2.63 | proton transmembrane transport |
|  | SMEL5_03g024080 | ABCG23 exporter | 2.71 | transmembrane transport |
|  | SMEL5_05g015090 | P450 71AU50-like | 2.84 | monooxygenase activity |
|  | SMEL5_04g005860 | FQR1-like NAD | 2.89 | NADPH dehydrogenase (quinone) activity |
|  | SMEL5_01g021840 | GST glutathione S-transferase | 2.89 | glutathione transferase activity |
|  | SMEL5_05g015160 | FAD2-like desaturase | 2.92 | unsaturated fatty acid biosynthetic process |
|  | SMEL5_05g015120 | FAD2-like desaturase | 2.96 | unsaturated fatty acid biosynthetic process |
|  | SMEL5_05g024330 | ADP/ATP carrier, mitochondrial | 3.24 | mitochondrial ADP transmembrane transport |
|  | SMEL5_02g027950 | PIOX-like alpha-dioxygenase | 4.04 | response to oxidative stress |
|  | SMEL5_04g015610 | UGT-like glucosyltransferase | 5.73 | negative regulation of fatty acid metabolic process |
|  | SMEL5_05g018500 | TUA3 tubulin alpha-3 | -1.03 | microtubule cytoskeleton organization |
|  | SMEL5_02g011300 | PSI subunit III | -1.06 | photosynthesis |
|  | SMEL5_06g028360 | PSI reaction center subunit XI | -1.21 | photosynthesis |
|  | SMEL5_10g020840 | TUBB2 tubulin beta-2 | -1.47 | structural constituent of cytoskeleton |
|  | SMEL5_05g007900 | Chlorophyll a-b binding protein 6A | -2.06 | photosynthesis, light harvesting in photosystem I |
|  | SMEL5_01g011400 | KIN-14I kinesin-like protein | -2.16 | oxidoreductase activity |
|  | SMEL5_01g007320 | CP24 10B | -2.24 | photosynthesis, light harvesting in photosystem I |
|  | SMEL5_07g022520 | KIN-12F kinesin-like protein | -2.34 | microtubule-based movement |
|  | SMEL5_05g007920 | Chlorophyll a-b binding protein 6A | -2.42 | photosynthesis, light harvesting in photosystem I |
|  | SMEL5_09g016950 | Cellulose synthase-like D5 | -2.44 | cellulose biosynthetic process |
|  | SMEL5_02g009780 | Chlorophyll a-b binding protein 21 | -2.48 | response to desiccation |
|  | SMEL5_04g010390 | Cyclin-A1-4-like | -2.76 | Cell division |
|  | SMEL5_09g023710 | KIN-12B kinesin-like protein | -2.77 | microtubule-based movement |
|  | SMEL5_02g009770 | Chlorophyll a-b binding protein 21 | -2.91 | response to desiccation |
|  | SMEL5_03g026860 | WIN1/SHN-like wax/cuticle regulator | -3.15 | triglyceride biosynthetic process |
|  | SMEL5_06g012380 | KNOLLE-like syntaxin | -3.57 | cell division |
|  | SMEL5_02g015680 | Pectinesterase-like | -4.21 | cell wall modification |
|  | SMEL5_04g004610 | Expansin-B3-like | -4.89 | Cell wall biogenesis/degradation |
|  | SMEL5_06g023140 | SWEET9-like sucrose efflux transporter | -5.08 | carbohydrate transport |
|  | SMEL5_06g023130 | SWEET12-like sucrose efflux transporter | -6.47 | carbohydrate transport |

Among suppressed biological processes at WS-M, enrichment was primarily associated with structural and growth-related functions, including cell wall organization or biogenesis, cutin-based cuticle development, and photosynthesis. Cuticle-related suppression was mirrored by downregulation of multiple epidermal barrier components, including an ABCG transporter (SMEL5_01g006970), CER3 (SMEL5_07g000910), a wax ester synthase/DGAT11-like enzyme (SMEL5_01g030260), a GPAT5-like gene (SMEL5_11g028180), and a strongly repressed MYB41-like regulator (SMEL5_04g005030; Table 1). Similarly, chloroplast- and photosynthesis-linked suppression was reflected by decreased expression of photoprotection and plastid maintenance nodes, including an ELIP1-like gene (SMEL5_09g018400), SOQ1 (SMEL5_01g010820/SMEL5_01g010830), a plastid RNA polymerase β′ (SMEL5_11g014820), and an NDH lumenal subunit (SMEL5_04g002050; Table 1). Water-transport categories were also negatively enriched at WS-M, with repression of specific aquaporins (TIP-type RB7-5A, SMEL5_06g014400; NIP2-1-like, SMEL5_05g012270) despite induction of other isoforms (e.g., TIP1-1-like, SMEL5_10g022180; Table 1).

GSEA confirmed a stage-specific functional redistribution at WS-S (Figure 4A,B). Negatively enriched categories were dominated by chlorophyll/pigment binding and photosynthetic light harvesting, mirrored by high-amplitude repression of multiple antenna and PSI loci, including chlorophyll a–b binding protein 21 (SMEL5_02g009770, SMEL5_02g009780), chlorophyll a–b binding protein 6A (SMEL5_05g007920, SMEL5_05g007900), CP24 10B (SMEL5_01g007320), and PSI components such as subunit XI (SMEL5_06g028360) and subunit III (SMEL5_02g011300; Table 1). In parallel, strong negative enrichment for microtubule cytoskeleton-, kinesin complex- and cell cycle–related terms was supported by repression of kinesins (KIN-12B, SMEL5_09g023710; KIN-12F, SMEL5_07g022520; KIN-14I, SMEL5_01g011400), a cyclin-A1-4-like gene (SMEL5_04g010390), tubulins (TUBB2, SMEL5_10g020840; TUA3, SMEL5_05g018500), and a KNOLLE-like syntaxin (SMEL5_06g012380; Table 1). Carbon allocation and tissue remodeling were concurrently affected, exemplified by strong repression of sucrose efflux transporters (SWEET12-like, SMEL5_06g023130; SWEET9-like, SMEL5_06g023140) and wall-associated loci such as expansin-B3-like (SMEL5_04g004610), a pectinesterase-like gene (SMEL5_02g015680), and cellulose synthase-like D5 (SMEL5_09g016950; Table 1).

Conversely, positively enriched gene sets at WS-S were dominated by mitochondrial and maintenance-associated functions, including respirasome/respiratory chain/oxidative phosphorylation/ATP synthesis, supported by induction of an ADP/ATP carrier (SMEL5_05g024330), a NADH–quinone oxidoreductase subunit (SMEL5_03g032040), and an FQR1-like NAD(P)H dehydrogenase (SMEL5_04g005860; Table 1). Severe drought also intensified membrane and chemical stress–associated programs, including lipid remodeling and oxylipin-linked enzymes (PIOX-like, SMEL5_02g027950; FAD2-like desaturases, SMEL5_05g015120, SMEL5_05g015160; PLIP1, SMEL5_01g015930; PLA1-II-like, SMEL5_02g006090), together with detoxification and redox-related components (GST, SMEL5_01g021840; metallothionein-like, SMEL5_04g010990) and conjugation/specialized metabolism enzymes (P450 71AU50-like, SMEL5_05g015090; UGT-like, SMEL5_04g015610; Table 1). Multiple exporters were also induced (ABCG23, SMEL5_03g024080; ABCG10-like, SMEL5_07g019280; PDR1-like, SMEL5_09g022110). Notably, within cuticle-related regulation, a WIN1/SHN-like wax/cuticle regulator (SMEL5_03g026860) was strongly repressed despite induction of ABC/PDR transporters and lipid-transfer associated genes (Table 1).

### Tolerant genotype shows ABA-centred early protection and controlled acclimation under increasing drought severity

In the tolerant genotype GPE020510, drought stress induced distinct and stage-dependent transcriptional changes during both the transition from well-watered conditions to moderate drought (T0 to WS-M) and from moderate to severe drought (WS-M to WS-S). At the moderate drought stage (T0 to WS-M), GPE020510 displayed differential regulation of a specific subset of genes (5925 upregulated and 6807 downregulated, Figure 6C), suggesting rapid engagement of dehydration-responsive protection, transport homeostasis, and redox buffering.

**Figure 6.**
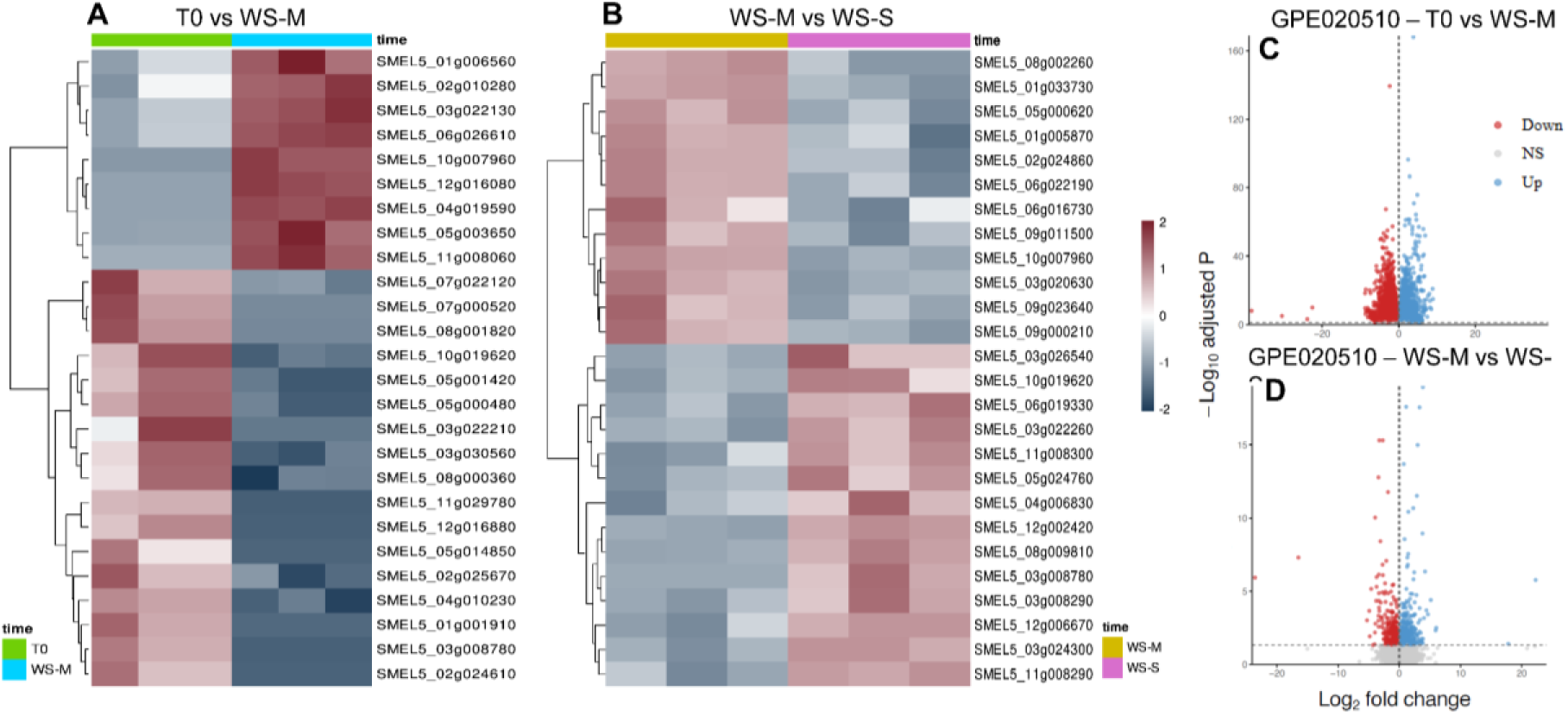
Stage-specific differential expression in response to progressive drought. Heatmaps (left) show scaled expression patterns of representative drought-responsive genes across samples, grouped by genotype and drought severity (A, T0 to WS-M; B, WS-M to WS-S; colour bars indicate sample classes). Genes are clustered by similarity, highlighting genotype- and stage-dependent transcriptional trajectories. Volcano plots (right) summarize differential expression for the corresponding contrasts (C, T0 vs WS-M; D, WS-M vs WS-S), with significantly upregulated genes shown in blue, downregulated genes in red, and non-significant genes in grey (dashed lines indicate the applied fold-change and adjusted P-value thresholds).

Among the most strongly induced transcripts were canonical ABA/dehydration regulators and effectors, including ABI5 (SMEL5_09g001060) and TAS14 (SMEL5_02g020190), together with prominent cellular protectants such as LEA/LEA2-like proteins (SMEL5_03g026640; SMEL5_01g002530) and Rab25-like dehydrin (SMEL5_02g003520) (Table 2). This ABA-centred response was accompanied by induction of additional stress-responsive regulators, including ATHB-7 (SMEL5_08g024810), a ZAT4-like zinc finger (SMEL5_01g019030), and an ERF2-like transcription factor (SMEL5_08g019370). In parallel, moderate drought in GPE020510 was characterized by prominent activation of transport and membrane homeostasis components, including a sodium-coupled neutral amino acid transporter (SMEL5_06g008410) and induced ABCG transporters (SMEL5_01g006930; SMEL5_11g019430), together with redox/detox-associated genes such as ETHE1 (SMEL5_10g004650) and a peroxygenase-like enzyme (SMEL5_05g021400), and a strongly induced low-temperature-induced 65 kDa protein-like transcript (SMEL5_03g015400) (Table 2). Concomitantly, the WS-M transition also included marked repression of growth-associated outputs, exemplified by multiple auxin-responsive SAUR68-like loci (SMEL5_11g009750; SMEL5_11g001820) and wall-associated genes such as a cellulose synthase-like locus (SMEL5_07g013090) and a polygalacturonase-like gene (SMEL5_03g010400) (Table 2). During the subsequent transition to severe drought (WS-M to WS-S), the transcriptional profile of GPE020510 shifted toward a broader involvement of stress acclimation and remodeling-related genes (742 upregulated and 556 downregulated, Figure 6D), with prominent signatures linked to plastid/light management and proteostasis. Induced photoprotection and light-handling nodes included ELIP1 (SMEL5_09g018430), OHP2 (SMEL5_05g017260), and BIC1/BIC1-like regulators (SMEL5_12g014390; SMEL5_12g014380) (Table 2). Severe drought was also associated with activation of dehydration/cold-related regulators and chaperone capacity, including DREB1A-like (SMEL5_03g014860), CBF/AP2 (SMEL5_03g014870), and small heat shock proteins (SMEL5_12g006670; SMEL5_03g017990) (Table 2). In parallel, GPE020510 showed induction of genes compatible with targeted metabolic adjustment, including carbohydrate- and lipid-associated drivers such as β-amylase 3 (SMEL5_08g000360), 6-phosphofructokinase 6-like (SMEL5_08g011260), phospholipase A1-II 1-like (SMEL5_02g006090), and a FAD2 desaturase (SMEL5_05g024580), alongside stress-linked oxidative and specialized metabolism enzymes including F6H1-3 (SMEL5_05g024760), premnaspirodiene oxygenase (SMEL5_11g013170), CYP94B3-like/CYP98A2-like (SMEL5_03g021570), and SMEL5_06g017490 (Table 2). Notably, this late phase also included selective repression of specific nodes, including PLIP1 (SMEL5_01g015930), KAT3 (SMEL5_08g007330), and a WRKY70-like factor (SMEL5_10g019970) (Table 2).

**Table 2.**
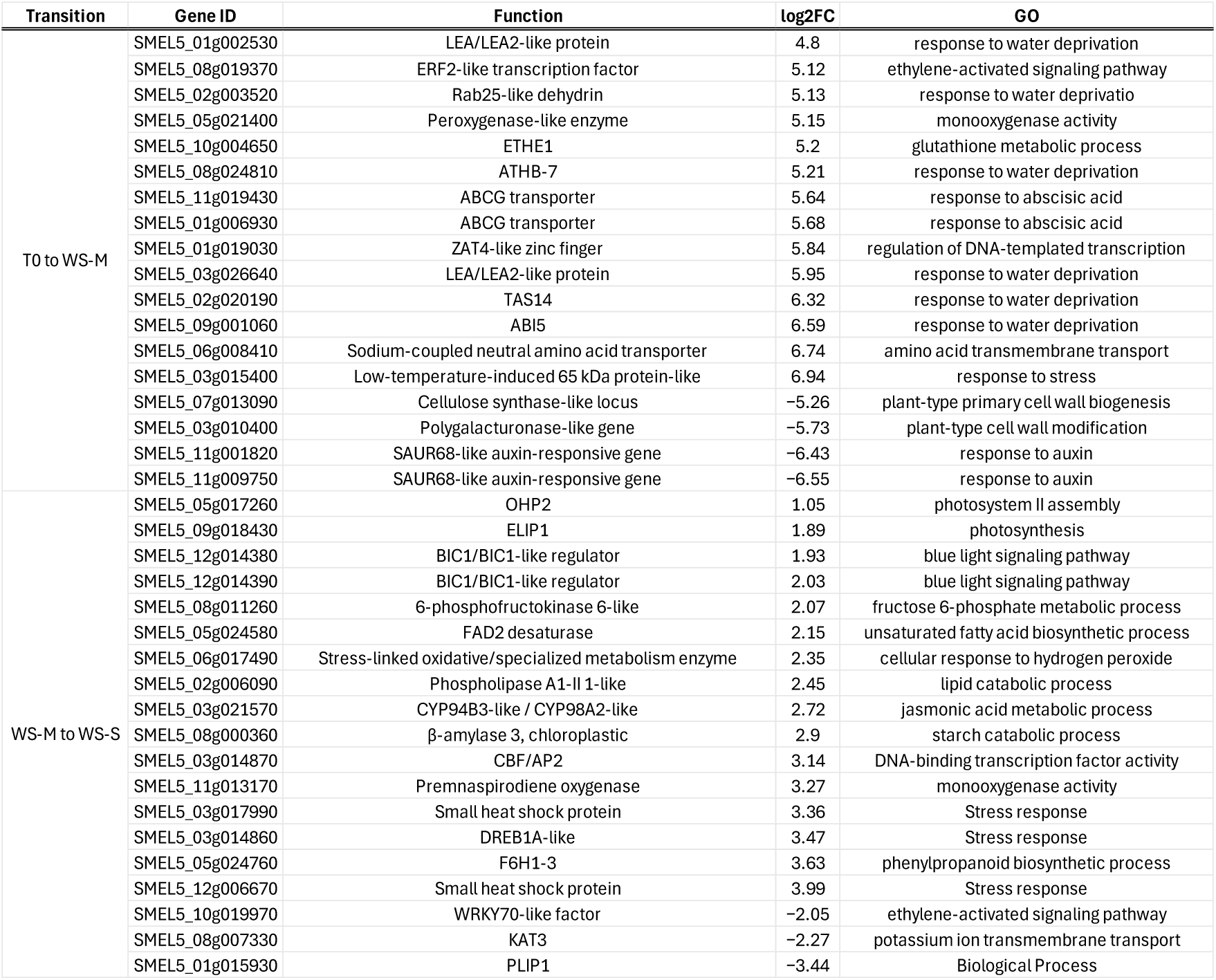
Key drought-responsive genes prioritized in the tolerant genotype GPE020510 across progressive drought severity. The table reports the selected candidate transcripts (gene IDs) highlighted in the Results for the “T0 to WS-M” and “WS-M to WS-S” transitions, with functional annotation, direction of regulation in each contrast, and gene ontology.

Gene set enrichment analysis of differentially expressed genes revealed stage-dependent functional enrichment patterns during the progression from well-watered conditions to moderate (WS-M) and severe drought (WS-S - Figure 7). At the early drought stage (T0 to WS-M), enrichment profiles showed a clear separation between gene sets associated with up-regulated and down-regulated transcripts. Within the Biological Process ontology at WS-M, up-regulated gene sets were predominantly associated with catabolic and respiratory processes, including carboxylic and organic acid catabolism, amino acid catabolism, small molecule catabolic process, and cellular respiration, together with regulation of proteolysis, lipid catabolic processes, and ribosomal small subunit biogenesis. In contrast, down-regulated biological processes were mainly related to photosynthesis and structural components, including cell wall organization or biogenesis, cutin-based cuticle development, water transport, and auxin polar transport. Consistent patterns were observed across the Molecular Function and Cellular Component ontologies. At WS-M, up-regulated gene sets were enriched for ion binding and transmembrane transporter activities, together with DNA- and protein-related catalytic functions, whereas down-regulated molecular functions included chlorophyll and pigment binding. At the cellular level, up-regulated gene sets were associated with protein- and degradation-related complexes, including proteasome- and spliceosome-associated components, while down-regulated gene sets were enriched for plastid- and photosynthesis-related compartments such as thylakoid membranes and plastoglobules.

**Figure 7.**
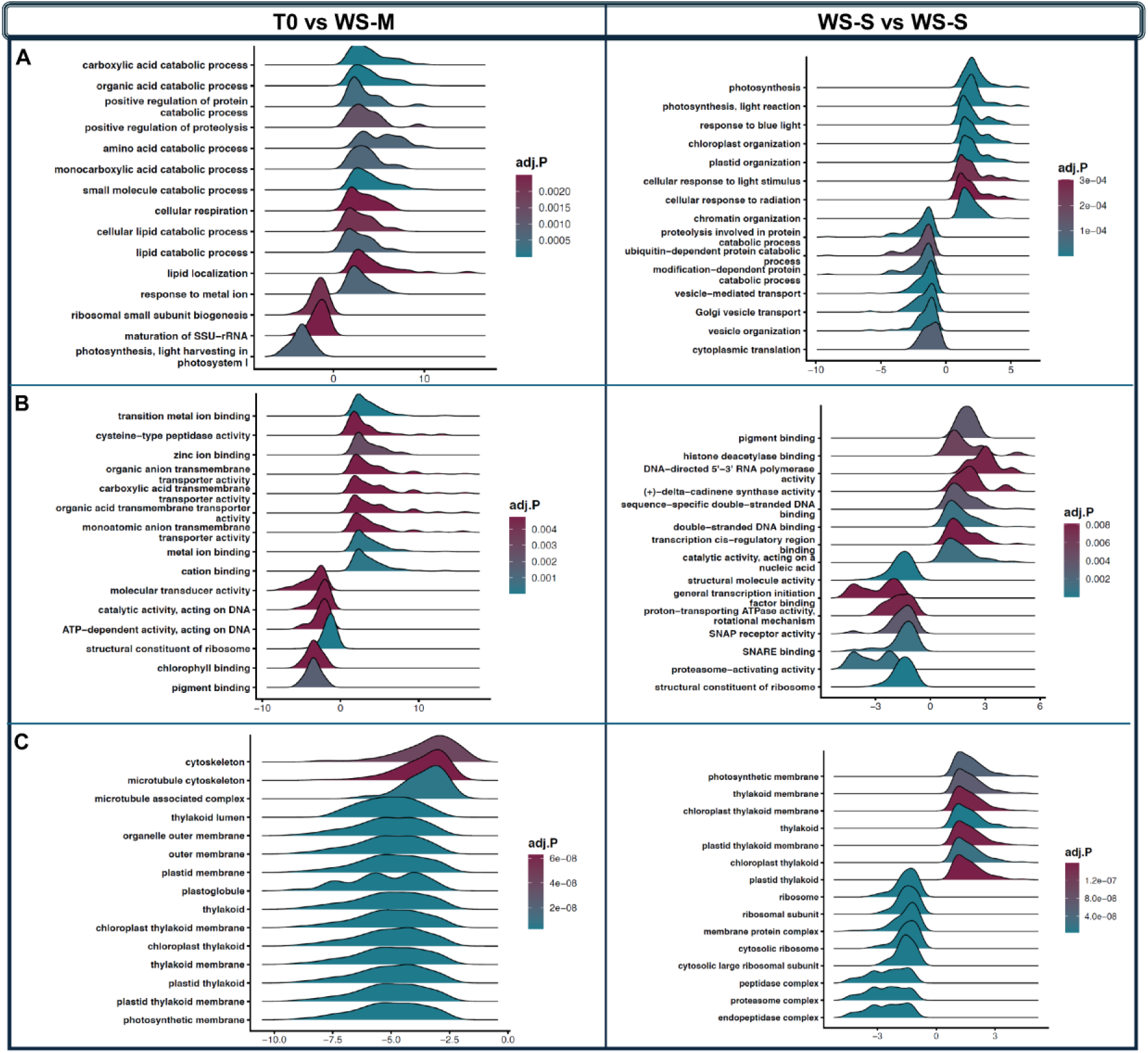
Ridge plots of gene set enrichment results across progressive drought in the tolerant genotype. Enriched terms are shown for Biological Process (top row), Molecular Function (middle row) and Cellular Component (bottom row). The left column summarizes the transition from well-watered to moderate drought (T0 to WS-M), while the right column summarizes the transition from moderate to severe drought (WS-M to WS-S). For each panel, ridges represent individual significantly enriched terms, positioned along the enrichment score axis (positive values indicate enrichment among upregulated gene sets; negative values indicate enrichment among downregulated gene sets). Ridge colour encodes the adjusted P value (adj. P).

### Co-expression modules associated with moderate-to-severe drought transition

Comparison of differentially expressed genes between the two genotypes at comparable drought stages revealed a very limited overlap. To further characterize transcriptional changes associated with the transition from T0 to severe drought, we performed a weighted gene co-expression network analysis (WGCNA) using samples collected at comparable physiological stages. This approach was employed to identify groups of co-expressed genes whose coordinated behavior differed between genotypes during the progression to severe water deficit (Figure 8).

**Figure 8.**
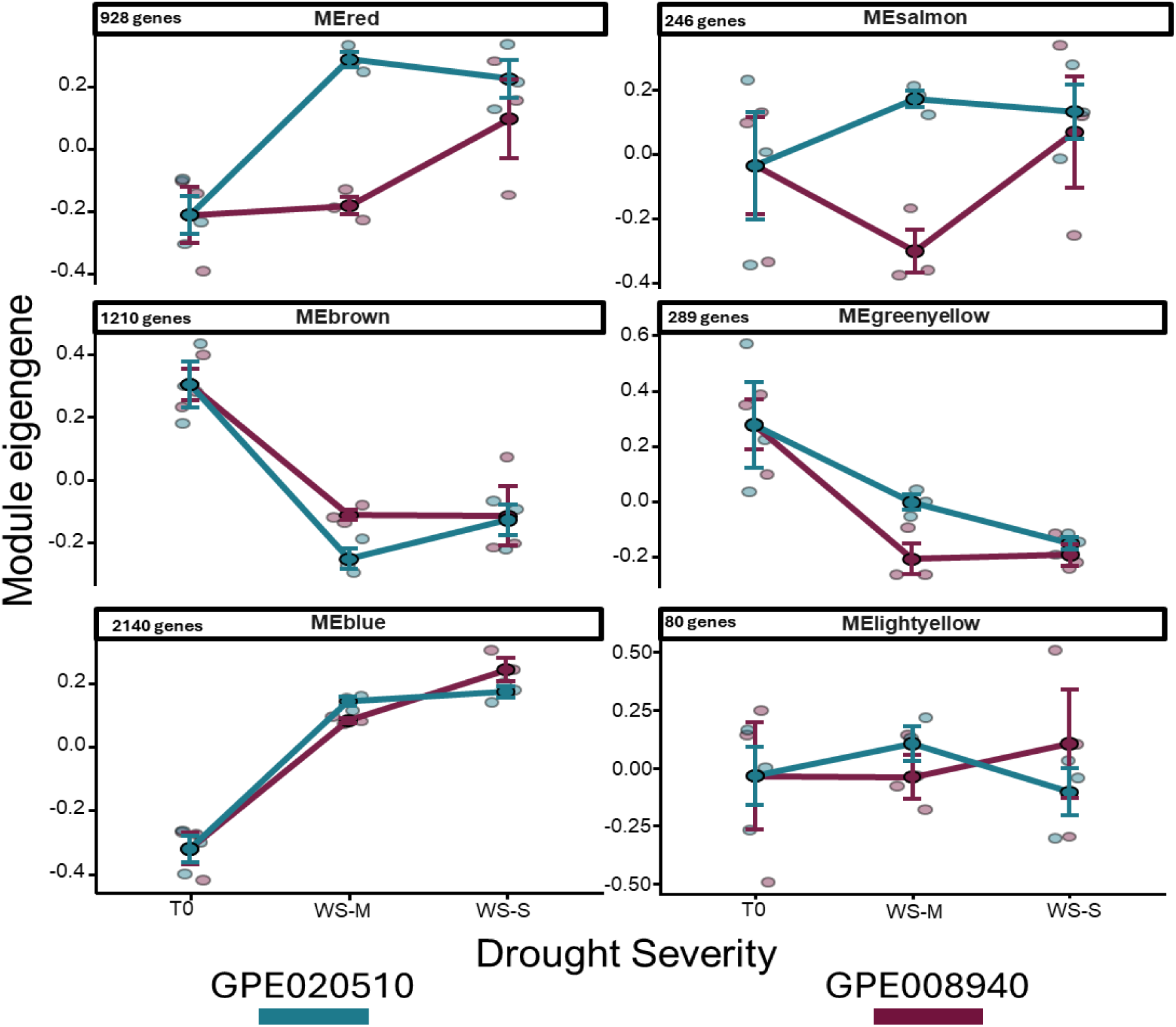
Eigengene trajectories of six WGCNA modules across increasing drought severity (T0 to WS-S) in the two genotypes. For each module, individual sample eigengene values are shown as points, and group means ± error are indicated by larger points with error bars, and genes per module are reported.

Six co-expression modules showed distinct eigengene trajectories across drought progression and differed between genotypes (Figure 8). Hub genes are listed in Supplementary Table 1. The red module increased strongly from T0 to WS-M in the tolerant genotype GPE020510 and remained high at WS-S, whereas in the susceptible genotype GPE008940 increased mainly at WS-S. Based on its hubs, this module is consistent with a regulatory/signaling timing program. Hub genes included SMEL5_05g017420 (WRKY51-like), SMEL5_06g018620 (ELF3-like), SMEL5_09g002410 (zinc finger 593-like), and SMEL5_02g020630 (epoxide hydrolase 4-like). The salmon module diverged between genotypes at WS-M: GPE008940 strongly decreased at WS-M and recovered by WS-S, whereas GPE020510 slightly increased at WS-M and remained positive at WS-S. Its hubs point to a membrane trafficking and secondary-metabolism/stress-marker program. Hub genes included SMEL5_01g036930 (syntaxin-121-like Qa-SNARE), SMEL5_05g015090 (P450 71AU50-like), SMEL5_01g004620 (scopoletin glucosyltransferase-like UGT), and SMEL5_06g000430 (SAG21-like). The brown module decreased sharply from T0 to WS-M in both genotypes and remained low at WS-S, with a stronger drop at WS-M in the tolerant genotype GPE020510 and partial recovery by WS-S. Its hub composition points to a chloroplast redox / tetrapyrrole /photosynthetic adjustment program. Representative hub genes included SMEL5_09g000310 (porphobilinogen deaminase), SMEL5_10g025240 (NTRC), and SMEL5_12g004410 (NDH lumenal subunit). The greenyellow module showed a strong reduction from T0 to WS-M followed by persistently low values at WS-S, with a more pronounced WS-M decrease in the susceptible genotype GPE008940 than in the tolerant genotype GPE020510. Its hubs suggest a PRR-like receptor kinases and Ca²⁺-linked signaling program. Hub genes included SMEL5_08g014220 and SMEL5_08g014230 (AED1-like aspartyl proteases), and receptor-kinase loci SMEL5_11g026670 (EFR-like) and SMEL5_11g025890 (Xa21-like). The blue module increased from T0 to WS-M in both genotypes and further increased toward WS-S, with higher WS-S values in the susceptible GPE008940 than in the tolerant GPE020510. Its hubs are consistent with a core escalation program involving ion/transport and RNA processing/signaling. Hub genes included SMEL5_05g010420 (plasma membrane H+-ATPase 1), SMEL5_01g011740 (SR30-like splicing factor), and SMEL5_10g012140 (VIP1-like). Finally, the lightyellow module displayed opposite trajectories between genotypes: the tolerant GPE020510 increased at WS-M and decreased at WS-S, whereas the susceptible GPE008940 remained near baseline at WS-M and increased at WS-S. Its hub genes support a transcription/translation and regulatory-complex program. Hub genes included SMEL5_12g003930 (SKP1-like 1B), SMEL5_06g024510 (RNA polymerase II subunit 4), SMEL5_02g014660 (eIF3J-A), and SMEL5_04g006610 (SOC1-like).

## Discussion

Physiological measurements conducted in this study (*i.e.*: stomatal conductance and leaf water potential, Fig. 1) confirmed the contrasting drought performance under progressive stress previously described within the G2P-SOL framework, with the genotype GPE020510 maintaining plant water status and stomatal function longer than the genotype GPE008940, suggesting improved drought tolerance (Asargew et al., 2024; Lu et al., 2020). At the transcriptome level, both genotypes showed strong reprogramming all over the period, with responses staged across WS-M and WS-S, defined for each genotype by comparable leaf water potential.

Under moderate stress (WS-M), the tolerant genotype GPE020510 was characterized by signatures points to early acclimation, including metabolic adjustment and ABA-linked regulatory control. This aligns with a well-established framework in which ABA-centred signaling and downstream effectors coordinate early dehydration responses that promote osmotic adjustment, cellular protection and homeostasis buffering (Liao et al., 2023; Soma et al., 2021; Wei et al., 2025). In our data, this early protective configuration is supported by candidate regulators/effectors such as ABI5 (SMEL5_09g001060) and TAS14 (SMEL5_02g020190), together with dehydration-associated protectants (*e.g.*, LEA/dehydrin-type loci; Table 2), core components of drought-responsive transcriptional cascades (Muñoz-Mayor et al., 2012; Skubacz et al., 2016; Vaseva et al., 2025). Importantly, the tolerant profile at WS-M also included growth restraint (*e.g.*, repression of SAUR-associated loci and wall-related genes), which is coherent with the broader concept that drought acclimation often requires prioritizing protection overgrowth through hormone-mediated reallocation (Jiang et al., 2025; Khan et al., 2023; Seleiman et al., 2021). In contrast, at WS-M the susceptible genotype GPE008940 showed a response biased toward regulatory rewiring, with enrichment for RNA/nuclear and turnover-associated functions, in line with reported transcript fate and protein composition changes under stress conditions (Irshad and Sharma, 2024). However, in the susceptible genotype GPE008940 this regulatory engagement coincided with an early downshift of two protective “infrastructure” layers: epidermal barrier/cuticle-associated programs and chloroplast/light-harvesting and plastid maintenance functions (Figures 4–5; Table 1). This is biologically meaningful because cuticle and wax pathways have been linked to limiting non-stomatal water loss and mitigating dehydration-associated injury (Chen et al., 2020; Han et al., 2026; Zhao et al., 2025), while chloroplast photoprotection and plastid gene-expression/maintenance are central to preventing excess excitation pressure and ROS build-up as CO₂ supply declines under drought (Didaran et al., 2024; Gao et al., 2026; Martins et al., 2023; S. Wang et al., 2025). Thus, the susceptible genotype appears to enter the trajectory toward severe stress with comparatively weaker deployment of barrier- and plastid-support outputs, plausibly increasing vulnerability to downstream homeostasis loss.

As drought intensified to WS-S, the susceptible genotype adopted a transcriptional configuration consistent with a low carbon gain/high maintenance state: strong repression of light harvesting and growth-related machinery together with increased representation of mitochondrial respiration, detoxification, membrane/lipid remodeling and transporter/export functions (Figures 4–5). This pattern matches widely described late drought physiology in which reduced photosynthetic productivity co-occurs with increased energetic and biochemical costs associated with damage control, redox management, and membrane maintenance (Chachar et al., n.d.; Chauhan et al., 2023; Nour et al., 2024; Qiao et al., 2024). By contrast, the tolerant genotype GPE020510 showed a more controlled severe-stress profile, with evidence of plastid/light-management and proteostasis-related adjustment and a more targeted metabolic response, mechanisms known for their role in stress regulations and mitigation (Cheng et al., 2025; Li et al., 2026; S. Wang et al., 2025; Wei et al., 2020). Together, the two-stage comparison supports the interpretation that susceptibility here reflects coordination and timing: early regulatory rewiring coupled to reduced barrier/plastid-support signatures, followed by stronger late maintenance/detox escalation, rather than a simple attenuation of the tolerant program.

WGCNA provided module-level support for the early/late phasing model (Figure 8), hub genes are listed in Supplementary Table X. A first axis involves regulatory/stress-signaling and membrane/trafficking modules that separate at WS-M. The red module rises early in the tolerant genotype GPE020510 and remains elevated, whereas the susceptible GPE008940 shows a stronger increase mainly at WS-S. Its hub composition is consistent with a coordinated stress-regulatory program integrating hormone-linked control, osmotic adjustment and redox/proteostasis components (Abbas et al., 2023; Ali et al., 2025; Jurado-Mañogil et al., 2024). In parallel, the salmon module highlights a membrane-centered branch of the response, with hubs including a SYP121/PEN1-class syntaxin (SMEL5_01g036930) together with lipid/secondary-metabolism and stress-marker nodes. Given the established role of SYP121/PEN1-class SNAREs in secretory traffic and guard-cell/stomatal function (Eisenach et al., 2012), these trajectories suggest that the tolerant GPE020510 engages trafficking-linked adjustment earlier, while the susceptible GPE008940 shows a delayed or differently phased deployment of the same module. A second axis emerging from the co-expression structure is plastid/redox-centered regulation. The brown module is enriched in chloroplast and redox/tetrapyrrole-linked hubs, including NTRC (SMEL5_10g025240), porphobilinogen deaminase (SMEL5_09g000310) and additional chloroplast redox and photoprotective components. The trajectory shows a pronounced shift already at WS-M (stronger in the tolerant genotype, with partial recovery toward WS-S), compatible with early tuning of chloroplast redox/energy metabolism and tetrapyrrole-linked processes (Karami et al., 2025; Nagahatenna et al., 2015; Suzuki et al., 2012; S. Wang et al., 2025), potentially limiting downstream ROS amplification during drought escalation. Beyond these core axes, the greenyellow module points to genotype-dependent modulation of a receptor/Ca²⁺/protease signaling layer. Its stronger downshift in GPE008940 suggests differential prioritization of this signaling program during stress entry, consistent with drought–defense cross-talk and resource-allocation trade-offs (Dwivedi et al., 2021; Leisner et al., 2023). In addition to these core axes, two modules capture broader reprogramming layers that become more apparent with drought escalation. The blue module shows a progressive increase from T0 to WS-M and further toward WS-S in both genotypes but reaches higher values under WS-S in the susceptible GPE008940. Its hub composition suggests a coordinated “core drought-coping” program integrating osmotic/ABA-linked responses, ion and membrane energization, redox buffering, and proteostasis/autophagy (Bandurska, 2022; De Rossi et al., 2021; Dreyer et al., 2024; Haghpanah et al., 2024; Muhammad Aslam et al., 2022; Refaiy et al., 2026; M. Wang et al., 2025; Yu et al., 2025). In contrast, the lightyellow module displays opposite temporal phasing between genotypes: it increases at WS-M and then declines toward WS-S in the tolerant GPE020510, whereas the susceptible GPE008940 remains closer to baseline at WS-M and increases mainly at WS-S. Notably, lightyellow is a small module (80 genes), so functional interpretation should be carefully evaluated. This module points to a compact transcription/translation and regulatory-complex program whose deployment is earlier in the tolerant genotype but shifted toward late-stage induction in the susceptible genotype, in line with a stronger requirement for severe-stage reprogramming as drought reaches WS-S.

Integrating DEG/GSEA with module-level trajectories supports a model in which drought tolerance in GPE020510 relies on earlier, coordinated acclimation, particularly plastid/redox and membrane-regulatory tuning, thereby dampening the need for broad late-stage maintenance and remodeling responses that dominate in GPE008940 at WS-S.

## Conclusion

This study provides a stage-resolved transcriptomic framework for progressive drought in eggplant, showing that tolerance and susceptibility differ in the coordination of regulatory and organelle-/barrier-centered programs across transitions rather than in response magnitude alone. The resulting gene-set and co-expression architecture defines a prioritized set of candidate processes and hub nodes and, critically, anchors them to specific drought stages. A logical extension will be to integrate these stage-specific signatures into genetic mapping and molecular breeding pipelines (*e.g.* by intersecting candidate genes/modules with QTL/association signals, developing informative molecular markers, and deploying them in marker-assisted selection) while leveraging a compact set of complementary physiological and anatomical readouts (plastid/redox stability, carbon balance, barrier integrity) to support validation and refine prioritization across genetic backgrounds and environments.

## Author contributions

**M.M**: Writing – review & editing, Writing – original draft, Visualization, Methodology, Investigation, Formal analysis, Data curation, Conceptualization; **C.M**: Writing – review & editing, Writing – original draft, Visualization, Methodology, Investigation, Formal analysis, Data curation, Conceptualization; **A.M**: Writing – review & editing, Visualization, Conceptualization; **A.M.M**.: Writing – review & editing, Methodology, Formal analysis ; **L.B**: Writing – review & editing, Methodology, Investigation; **A.A**. : Writing – review & editing, Methodology, Investigation; **C.C**: Writing – review & editing, Methodology, Investigation, Conceptualization; **F.S**: Writing – review & editing, Methodology, Investigation, Conceptualization; **E.P**: Writing – review & editing, Methodology, Investigation, Conceptualization.

## Data availability

Sequencing data used in this study are openly available in the NCBI database (PRJ(.

